# Role of African swine fever virus (ASFV) proteins EP153R and EP402R in reducing viral persistence and virulence from attenuated BeninΔDP148R

**DOI:** 10.1101/2021.04.16.439957

**Authors:** Vlad Petrovan, Anusyah Rathakrishnan, Muneeb Islam, Lynnette C. Goatley, Katy Moffat, Pedro J. Sanchez-Cordon, Ana L. Reis, Linda K. Dixon

## Abstract

The limited knowledge on the role of many of the approximately 170 proteins encoded by African swine fever virus restricts progress towards vaccine development. In this study we investigated the effect of deleting combinations of different genes from a previously attenuated virus, BeninΔDP148R on: virus replication in macrophages, virus persistence and clinical signs post immunization, and induction of protection against challenge. Deletion of either EP402R or EP153R genes individually or in combination from BeninΔDP148R did not reduce virus replication *in vitro*. However, deletion of EP402R dramatically reduced viral persistence *in vivo*, whilst maintaining high levels of protection against challenge. The additional deletion of EP153R (BeninΔDP148RΔEP153RΔEP402R) further attenuated the virus and no viremia or clinical signs were observed post immunization. This was associated with decreased protection and detection of moderate levels of challenge virus in blood. Interestingly, the deletion of EP153R alone from BeninΔDP148R did not result in further virus attenuation and a slight increase in virus genome copies in blood was observed at different times post immunization when compared with BeninΔDP148R. These results show that EP402R and EP153R have a synergistic role in promoting viremia, however EP153R alone does not seem to have a major impact on virus levels in blood.

## Introduction

African swine fever virus (ASFV) causes an acute hemorrhagic fever in domestic pigs and wild boar with case fatality approaching 100%. In contrast, its wildlife hosts in Africa, warthogs, bushpigs and soft ticks of *Ornithodoros* species that inhabit warthog burrows can be persistently infected, but show few disease signs (1, 2). ASF has a very high economic impact in affected countries, which now include most sub-Saharan Africa, parts of Russia, Eastern Europe and 11 EU countries. The spread to China in 2018 and subsequent spread to South-East Asia resulted in death or culling of more than 7 million pigs and a decrease of the Chinese herd by about 40% (http://www.fao.org/ag/againfo/programmes/en/empres/ASF/situation_update.html). Since there are no vaccines or targeted therapeutics currently available, control relies on implementing strict biosafety and biosecurity measures.

ASFV is a large DNA virus with a linear double-stranded genome varying in size from 170 to more than 190 kbp. The virus is the only member of the *Asfarviridae* family and has a predominantly cytoplasmic replication. The virus genome contains up to 167 genes including many that are not required for the virus to replicate in cells, but have roles in interactions with the host, to facilitate its survival and transmission (3). For example, the virus codes for several proteins that help the virus to evade the host innate immune response such as the type I interferon (IFN) response and apoptosis (4). Deletion of genes that inhibit type I IFN response, including members of the multigene families (MGF) 360 and 505 and DP96R (also designated UK) (5-10) can reduce the virulence of the virus in pigs, and induce an immune response to protect the animal against lethal challenge with a related virulent virus.

ASFV also codes for two transmembrane glycoproteins that are not essential for virus replication in cells, pEP402R (CD2v) and EP153R (11, 12). The EP402R gene codes for a type I transmembrane protein with similarity in its extracellular domain to the host CD2 protein. This virus protein pEP402R, also designated CD2v or CD2-like, is required for the binding of red blood cells to infected macrophages (haemadsorption or HAD) (13, 14). It is also presumed to cause binding of red blood cells to extracellular virions, as 95% of virus in blood from infected pigs was shown to be in the red blood cell fraction (15). The CD2v protein was also suggested to have a role in the ASFV induced inhibition of *in vitro* proliferation of lymphocytes in response to mitogens since deletion of the gene abrogated this effect (16). Interactions of proteins including SH3P7/mAbp1 and AP1 with the cytoplasmic tail were demonstrated, suggesting that these may be involved in intracellular trafficking of the protein through the Golgi apparatus (17, 18). The CD2v protein is the only virus protein to be detected on the surface of extracellular virions (19). Interestingly, pigs immunized with a recombinant CD2v, expressed in baculovirus, showed reduced viremia after challenge with E75 isolate (20). Moreover, an involvement of cell mediated protection was suggested, since several T-cell epitopes were mapped using overlapping CD2v peptides (21).

The EP153R type II transmembrane protein contains a predicted C-type lectin domain. C-type lectins are Ca2+-dependent glycan-binding proteins that are involved in cell-cell adhesion playing key roles in both innate and adaptive immune responses. For example, C-Type Lectin Receptors (CLRs) are important for recognition and capture of pathogens as these pattern recognition receptors (PRRs) have a high affinity for their ligands, which results in internalization of the pathogens (22). The EP153R protein has been demonstrated to augment the HAD induced by the CD2v protein (23) and to have roles in inhibiting apoptosis mediated by the p53 pathway and in reducing the surface expression of swine leucocyte antigen I (SLAI) (24, 25).

Our previous experiments showed that deletion of the DP148R gene from the genome of the virulent Benin 97/1 genotype I isolate, attenuated the virus in pigs and induced high levels of protection against lethal challenge with parental virus (7). In pigs immunized with BeninΔDP148R, a peak of viremia was detected in blood at 5- or 6-days post-infection, coincident with clinical signs. After this period, clinical signs were not observed, but virus genome in blood declined slowly over a period of about 60 days and infectious virus declined more rapidly and was not detected by about day 30 (7).

In our current study, we investigated the hypothesis that virus binding to red blood cells may prolong the persistence of virus in blood. Therefore, we deleted either EP402R, EP153R genes alone or in combination from the genome of the BeninΔDP148R, and carried out immunization and challenge experiments in pigs. The results confirmed that deleting the EP402R gene reduced virus persistence in blood. Transient clinical signs and a peak of viremia were still observed post-immunization of pigs and high levels of protection were observed. Deletion of the EP153R gene from the BeninΔDP148RΔEP402R genome further attenuated the virus and no clinical signs or viremia were observed. However, deletion of EP153R from BeninΔDP148R revealed a similar clinical outcome compared with BeninΔDP148R. Interestingly levels of virus in blood were higher in BeninΔDP148RΔEP153R.

Overall, using a genetic backbone of a previously attenuated virus, we provided new insights into the role of the CD2v and EP153R proteins in virus virulence and persistence.

## Materials and methods

### >1. Viruses and cells

The ASFV Benin 97/1 wild type and BeninΔDP148R isolates have been described elsewhere (7, 26). Deletion mutant viruses were cultured in porcine bone marrow cells (PBM). Titration of wild type virus was carried out by haemadsorption assay (in which the results are presented as HAD_50_/ml) or end-point titers based on mNeonGreen or Tag-RFP-T markers and presented as TCID_50_/ml, calculated using Spearman and Karber formula.

### >2. Construction of recombinant BeninΔDP148RΔEP402R, BeninΔDP148RΔEP153RΔEP402R and BeninΔDP148RΔEP153R

#### >i. Transfer plasmids

Transfer plasmid pΔEP402R-VP72GUS was constructed by cloning amplified right and left regions of the Benin 97/1 flanking the EP402R gene into the previously published pLoxPVP72GUSLoxP vector (27).

Vector pLoxPVP30TagRFP-TLoxP was generated by first amplifying the TagRFP-T gene (28) with a forward primer containing *Bam*HI restriction site (italic) and the ASFV P30 promoter sequence (bold) (29) (5’-*GGATCC***TTATTATTTTATAATTTTAAAATTGAATGGATTTTATTTTAAATATATCC** ATGGTGTCTAAGGGCGAAGAGCT) and a reverse primer containing a ASFV transcription termination signal (bold) (30) and *Eco*RI restriction site (italic) (5’–*GAATTC***AAAAAAAAAA**CTTGTACAGCTCGTCCATGCCAT). The TagRFP-T fluorescent marker was kindly provided by Dr. Chris Netherton (The Pirbright Institute, UK). The amplified product was then swapped with the VP72GUS cassette of pLoxPVP72GUSLoxP vector. Using this newly produced pLoxPVP30TagRFP-TLoxP, the left and flanking regions of DP148R were then cloned to produce transfer plasmid pΔDP148R-VP30TagRFP-T.

The other transfer plasmids pΔEP153R-VP30mNG and pΔEP153RΔEP402R-VP30mNG were synthesized commercially (Genscript, US). Plasmid pΔEP153R-VP30mNG contains the left and right flanking regions of EP153R, while pΔEP153RΔEP402R-VP30mNG contains the left flanking region of EP153R and the right flanking region of EP402R. In both plasmids, between the flanking regions, a reporter gene, mNeonGreen (mNG) under control of the ASFV VP30 promoter flanked with LoxP sites was added (31).

#### >ii. Homologous recombination and virus purification

Recombinant ASFV BeninΔDP148RΔEP402R was produced in a sequential 2 step deletion method. First, BeninΔEP402R (Figure 1B) was produced by infecting primary porcine alveolar macrophages (PAM) with Benin 97/1 (Figure 1A) and then transfection with pΔEP402R-VP72GUS using the TransIT-LT1 transfection reagent (Mirus Bio, USA). In the presence of X-Gluc, recombinant viruses expressing the GUS gene were identified and purified by multiple rounds of limiting dilutions. Next, using purified BeninΔEP402R as parental virus, homologous recombination was undertaken with transfer plasmid pΔDP148R-VP30TagRFP-T in wild boar lung cells (WSL-R). Cells expressing the TagRFP-T were identified via fluorescence activated cell sorting (FACS); single cells were isolated and cultured in individual wells of 96 well plates containing PBMs. The recombinant virus was subsequently purified by FACS using the method described by Rathakrishnan et al. (31). Similarly, to produce recombinants BeninΔDP148RΔEP153R (Figure 1C) and BeninΔDP148RΔEP153RΔEP402R (Figure 1D), WSL-R cells were infected with the BeninΔDP148R virus at a multiplicity of infection (MOI) of 2 and transfected with the pΔEP153R-VP30mNG or pΔEP153RΔEP402R-VP30mNG plasmids respectively using the TransIT-LT1 transfection reagent. Expression of mNeonGreen marker was monitored and recombinant viruses were isolated and purified by FACS and limiting dilution as described (31). Sequencing across the site of deletion confirmed the expected deletion and site of reporter gene insertion.

**Figure 1.**
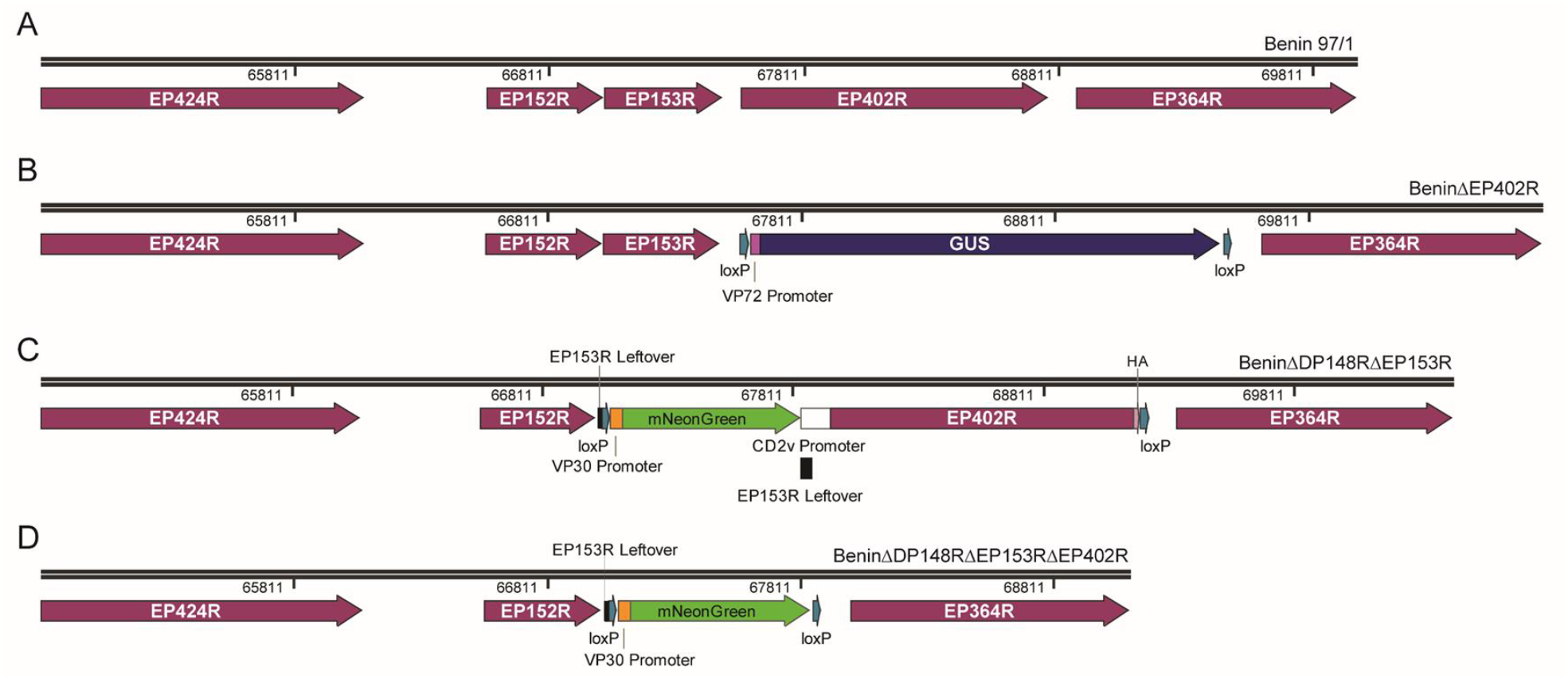
Schematic diagram depicting the deletion of EP153R and/or EP402R genes from genotype I ASFV. EP402R was deleted by homologous recombination between transfer plasmid pΔEP402R-VP72GUS and parental Benin 97/1 isolate **(A)** and the resultant, BeninΔEP402R **(B)**, was purified using limiting dilutions. Using a sequential deletion method, DP148R was then further deleted to produce BeninΔDP148RΔEP402R via single cell isolation, combined with limiting dilutions. With a previously described BeninΔDP148R, EP153R was deleted by homologous recombination to produce BeninΔDP148RΔEP153R **(C)**, which contains mNeonGreen reporter marker. Likewise, using BeninΔDP148R as the parental virus, a triple gene deleted virus, BeninΔDP148RΔEP153RΔEP402R **(D)** was produced. Both recombinant viruses were isolated and purified using FACS and limiting dilutions.

### >3. Growth curves

Viruses (Benin 97/1, BeninΔDP148R, BeninΔDP148RΔEP402R, BeninΔDP148RΔEP153RΔEP402R and BeninΔDP148RΔEP153R) were added to purified PBMs at a MOI of 0.01 in triplicate in 24 well plates. Cells and supernatants were collected at different times post-infection and subjected to 3 freeze-thaw cycles. Cellular debris was removed by centrifugation, and virus titers were determined by the fluorescence assay or HAD, as described above. The experiment was carried out in purified PBMs from 2 different pigs.

### >4. Immunization and challenge of pigs

Animal experiments were carried out in SAPO4 high containment animal housing at the Pirbright Institute according to regulated procedures from the Animals Act UK 1998 and conducted under Home Office License 7088520. Four separate experiments were undertaken. In experiment 1, 4 Large White X Landrace pigs (15-20kg) were immunized by the intramuscular route with 10^3^ TCID_50_ BeninΔDP148RΔEP402R and boosted by the same route with the same recombinant virus at 10^4^ TCID_50_ on day 20 post-immunization. In experiment 2, 8 pigs were immunized by the intramuscular route with 10^4^ TCID_50_ BeninΔDP148RΔEP153RΔEP402R and were boosted twice with the same virus at 10^4^ TCID_50_ on day 21 and at 10^6^ TCID_50_ on day 28 post-immunization. In experiment 3, 6 pigs were immunized by the intramuscular route with 10^5^ TCID_50_ BeninΔDP148RΔEP153R and boosted with the same dose on day 21 post-immunization. In experiment 4, 6 pigs were immunized by the intramuscular route with 10^5^ TCID_50_ BeninΔDP148R and boosted with the same dose on day 21 post-immunization. Non-immunized control pigs and immunized pigs were challenged by the intramuscular route with either 10^4^ (Exp. 1, group E) or 10^3^ (Exp. 2, group F, for Exp. 3, group L and for Exp. 4, group M) HAD_50_ of the virulent Benin 97/1 isolate approximately 3 weeks after the boost (Figure 2). Pigs were euthanized at a moderate severity endpoint as defined in the project license PPL70/8852 from the UK Home Office.

**Figure 2.**
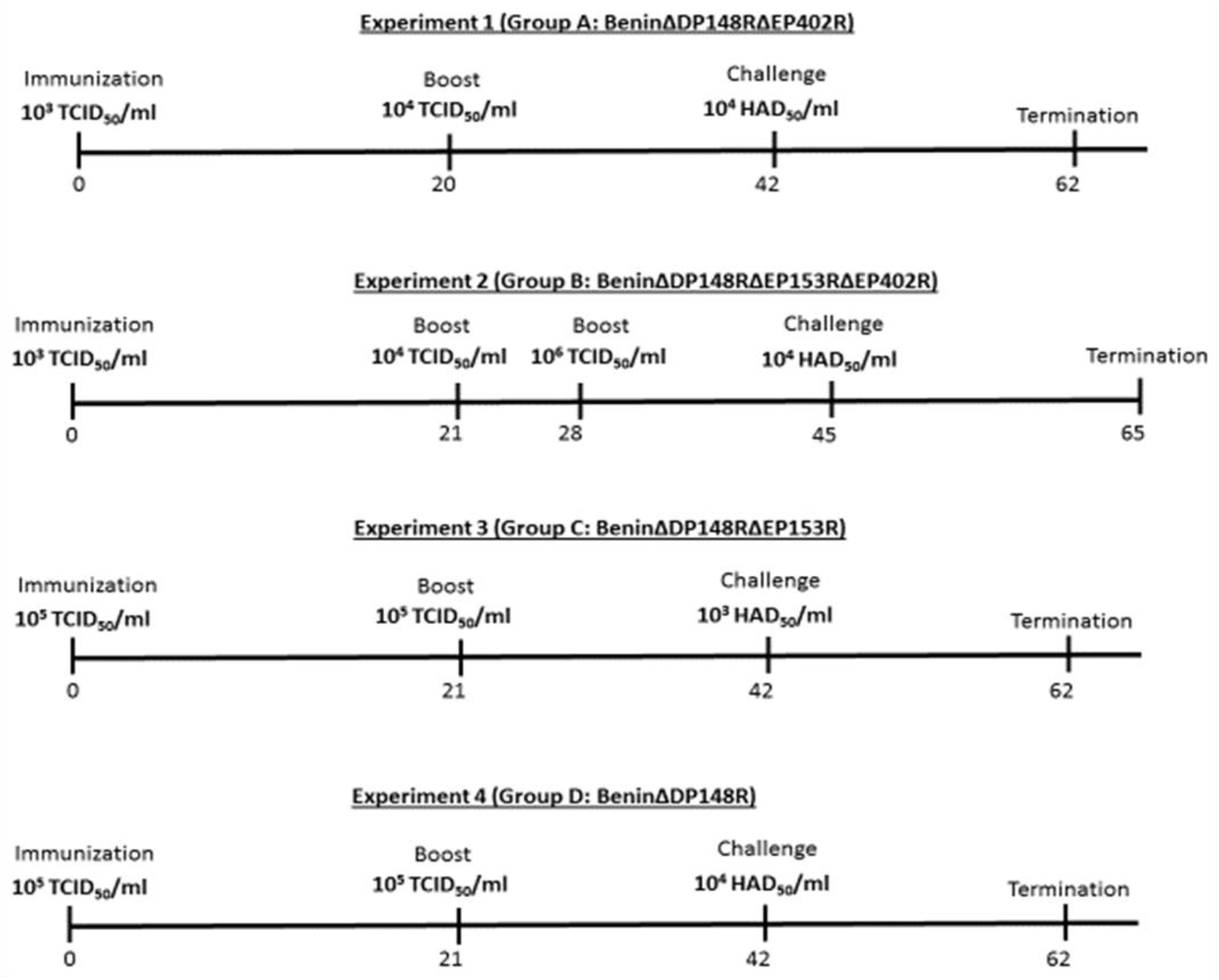
Timeline for the vaccination experiments. The days are given as post-immunization, beginning at day 0 with immunization, followed by boost, challenge with Benin 97/1 and finally termination. The amount of viruses used is given as either TCID_50_/mL or HAD_50_/mL

### >5. Genome copies in blood

DNA was extracted from whole blood using MagVet universal isolation kit (Life Technologies), at different days throughout the study. Samples were assayed in duplicate for the presence of viral DNA by qPCR on a Stratagene Mx3005P system (Agilent Technologies, Santa Clara, CA, USA) following a protocol modified (32) from using the primers Vp72 sense (CTG CTC ATG GTA TCA ATC TTA TCG A) and Vp72 antisense [GAT ACC ACA AGA TC(AG) GCC GT] and the probe 5′-(6-carboxyfluorescein [FAM])-CCA CGG GAG GAA TAC CAA CCC AGT G-3′-(6-carboxytetramethylrhodamine [TAMRA]) (32). A standard curve was prepared from a p72 mimic plasmid by making serial dilution ranging from 10^8^ to 10^1^ copies/ml. Results were reported as log10 genome copies/ml.

### >6. Antibody responses

The level of antibody responses against ASFV-p72 in serum was measured using a commercial blocking ELISA (INgezim PPA Compac, Ingenasa) following the manufacturer’s instructions. The percentage of blocking (PB) was calculated using the following formula: [(negative-control OD - sample OD) / (negative-control OD - positive-control OD)] × 100, where OD is optical density. Samples were considered positive if the PB was above the cut-off value of 50% blocking.

### >7. IFN gamma ELISpot assay

Peripheral blood mononuclear cells (PBMC) were collected at 3 different time points including pre-immunization, pre-boost and pre-challenge. PBMC were purified from EDTA-blood tubes using Histopaque-1083 gradient medium. ELISpot plates were coated overnight at 4°C with 4 μg/ml IFNγ monoclonal antibody (P2F6) in 0.05M carbonate-bicarbonate coating buffer. After incubation, plates were washed four times with phosphate-buffered saline (PBS). Cells were plated in duplicate at two different dilutions (8 × 10^5^ and 4 × 10^5^ per well), in RPMI 1640, Glutamax (Gibco) supplemented with 10% fetal bovine serum, 50 μM 2-mercaptoethanol, 100 IU/ml penicillin, and 100 μg/ml streptomycin. The cells were then incubated overnight at 37°C in a final volume of 200 μl with 10^5^ HAD_50_ of Benin 97/1, an equivalent volume of mock inoculum, or 20 μg/ml phytohemagglutinin as a positive control. Cells were lysed by incubation for 5 min in water and then washed with PBS. Following incubations with biotinylated anti-porcine IFNγ monoclonal antibody (P2C11) and streptavidin conjugated to alkaline phosphatase, AP conjugate substrate kit (Bio-Rad) was used to develop spots. The spot forming cells were then counted using an ELISpot assay reader system (Immunospot, CTL). The number of spots per well was converted into the number of spots per million cells, and the mean for duplicate wells was plotted using GraphPad Prism 8 software. No cells were collected for pigs belonging to group D (BeninΔDP148R).

### >8. Statistical analysis

Statistical analysis was performed using GraphPad Prism8 software. Two-way ANOVA followed by Sidak’s multiple comparison test was used to evaluate differences between groups.

## Results

### Generation of recombinant viruses

#### >i BeninΔDP148RΔEP402R

A two-step sequential deletion method was applied to produce the recombinant ASFV BeninΔDP148RΔEP402R (Figure 1B). In the first step, EP402R (Genome position: 67567 – 68775) was deleted from the virulent Benin 97/1 isolate (Figure 1A) using homologous recombination. This virus, BeninΔEP402R was subsequently cultured and used in the second step to construct recombinant ASFV BeninΔDP148RΔEP402R.

Using a single cell isolation method for producing recombinant ASFV (31), the second step in the production of BeninΔDP148RΔEP402R involved the inclusion of a fluorescent reporter gene TagRFP-T in place of the deleted DP148R gene. Using FACS, single cells expressing the TagRFP-T reporter were isolated and the recombinant virus was purified with a combination of single cell isolation and limiting dilutions.

#### >ii. BeninΔDP148RΔEP153RΔEP402R and BeninΔDP148RΔEP153R

A similar approach of single infected cell isolation and purification was used to generate recombinant ASFV BeninΔDP148RΔEP153R and BeninΔDP148RΔEP153RΔEP402R in which the either EP153R (genome position: 67051 - 67491) alone was deleted from BeninΔDP148R or both genes EP153R and EP402R (genome position: 67051 – 68775) were deleted simultaneously. These gene(s) were replaced by mNeonGreen under the control of VP30 promoter in the attenuated BeninΔDP148R virus (7) (Figures 1C and 1D). In both viruses, 21bp at the 5’ end of EP153R was left in the recombinant viruses because this stretch of sequence may contain the termination signal for the adjacent EP152R gene (33). The expected deletions of genes in the recombinant virus were confirmed via PCR analysis and Sanger sequencing. The purified recombinant virus stock was propagated on PBMs and titrations were performed in quadruplicate on PBMs collected from different pigs.

### Growth curves

Porcine bone marrow cells were infected with the Benin 97/1, BeninΔDP148R, BeninΔDP148RΔEP402R, BeninΔDP148RΔEP153RΔEP402R and BeninΔDP148RΔEP153R at a MOI of 0.01 to determine if deletion of the genes affected the ability of the virus to replicate *in vitro*. At different days post infection (1, 2, 3, 4, 5), total virus harvested from cells and supernatant was titrated. The results showed no significant difference between the kinetics and the levels of virus replication of the recombinant viruses and parental Benin 97/1 isolate (Figure 3). Virus titers reached a plateau of approximately 10^7^ TCID_50_/ml between 24 and 48 h post infection and were maintained for the remainder of the culture time. The results show that deletion of the DP148R, EP153R, EP402R or EP153R and EP402R gene did not significantly alter the ability of the virus to replicate in culture.

**Figure 3.**
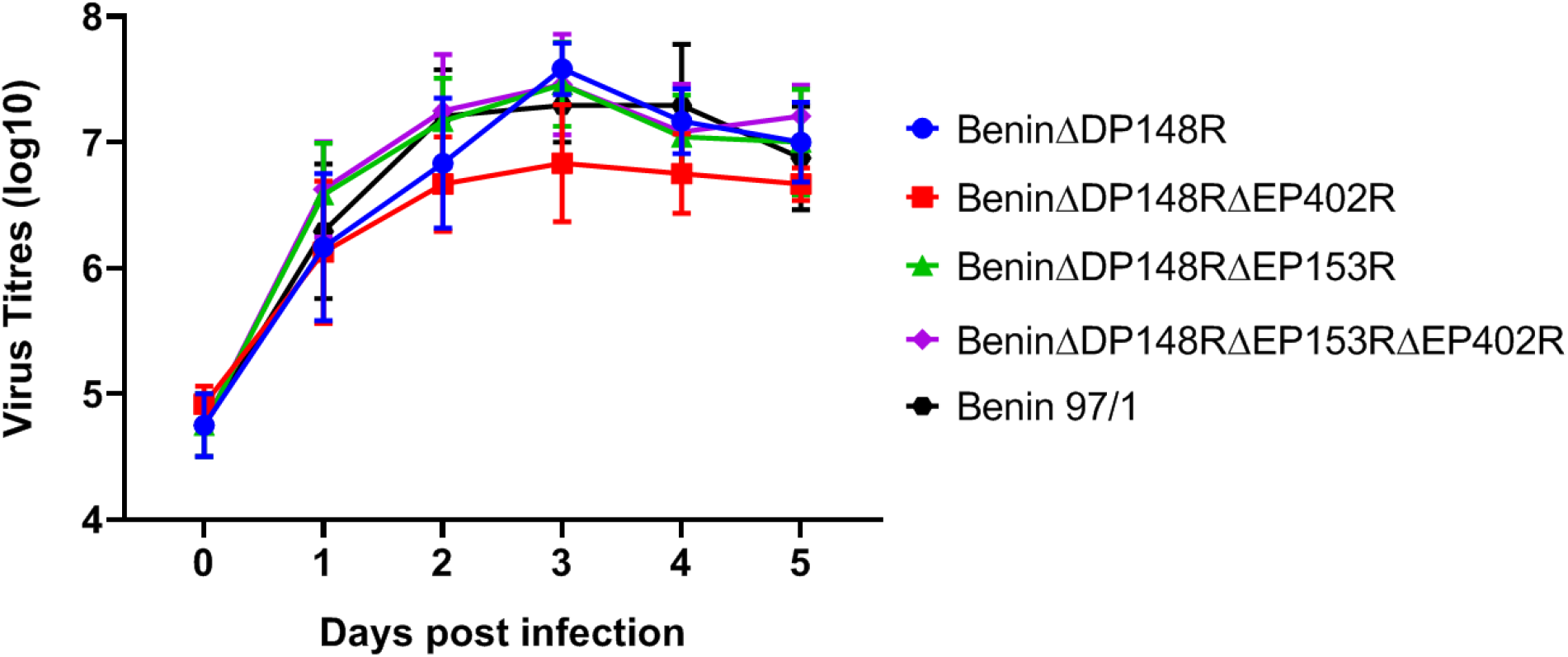
Replication of the recombinant gene deleted ASFV viruses compared to the wildtype Benin 97/1 strain. Purified PBMs from 2 different pigs were infected with viruses at MOI 0.01 in triplicates. Viruses were harvested from both cells and supernatants at different time points and titrated on PBMs in quadruplicates. Virus titres are presented as log10 HAD_50_/ml or TCID_50_/ml.

### Immunization, challenge and clinical observations

Four groups of pigs were immunized intra-muscularly (IM) in separate experiments with 1 ml of either 10^3^ TCID_50_ BeninΔDP148RΔEP402R (Group A), 10^4^ TCID_50_ BeninΔDP148RΔEP153RΔEP402R (Group B), 10^5^ TCID_50_ BeninΔDP148RΔEP153R (Group C) and 10^5^ TCID_50_ BeninΔDP148R (Group D). Pigs in all four groups were boosted with the same recombinant virus (with either 10^4^ for groups A and B or 10^5^ for groups C and D) approximately 3 weeks post immunization (see Figure 2). In Group B, an additional boost with 10^6^ TCID_50_ was performed as measurement of cellular responses at day 21 indicated low levels of ASFV specific IFNγ producing cells had been induced (see Figure 9). Naïve, non-vaccinated pigs, (Groups E, F, L and M) served as controls for the challenge with the virulent Benin 97/1 isolate. Groups A and E were challenged intramuscularly at 42 dpi with 10^4^ HAD_50_ in 1 ml while Group B and Group F were challenged at 45 dpi with 10^3^ HAD_50_ in 1 ml with virulent Benin 97/1 virus (Figure 2). One pig in Group A was euthanized at day 9 post-immunization due to a non-ASFV specific condition. Groups C and L were challenged at 39 dpi, whereas groups D and M were challenged at 42 dpi.

Rectal temperatures and clinical scores (34) were recorded daily for all pigs (Figures 4 and 5). Pigs in Group A (BeninΔDP148RΔEP402R) had transient increased temperatures above 40.5°C for 2 days after day 5 post-immunization (Figure 4A). This was accompanied by reduced appetite and lethargy (Figure 5A). Pig A1 had a temperature between 40 and 40.5 for 3 days (6, 7 and 9) and one day above 41 (day 8 - 41.2). A2 had a temperature at 41 or above for 2 days (6 and 8) and a temperature 40.3 on day 7. Pig A3 had a temperature of 40.6 on day 5 and 40.0 on day 6. One pig, A4, had a temperature of 40.4 on day 6, 40.8 on days 7 and 8 and 40.0 on day 9. This pig vomited blood and was euthanized on day 9 post-immunization. Post-mortem examination showed the pig had a stomach ulcer which was not suspected to be directly related to ASFV infection. No further clinical signs were observed post-immunization in the remaining pigs even after challenge. As expected, the non-immunized control pigs in Group E developed clinical signs associated with acute ASF after challenge. These signs included an increase in temperature (40.6 – 41.6°C), not eating and lethargy on day 3 post-challenge (Figures 4A and 5A). Pig E2 was also vomiting on day 4 post-challenge. All 3 pigs were culled on day 4 post-challenge at the moderate severity humane endpoint.

**Figure 4.**
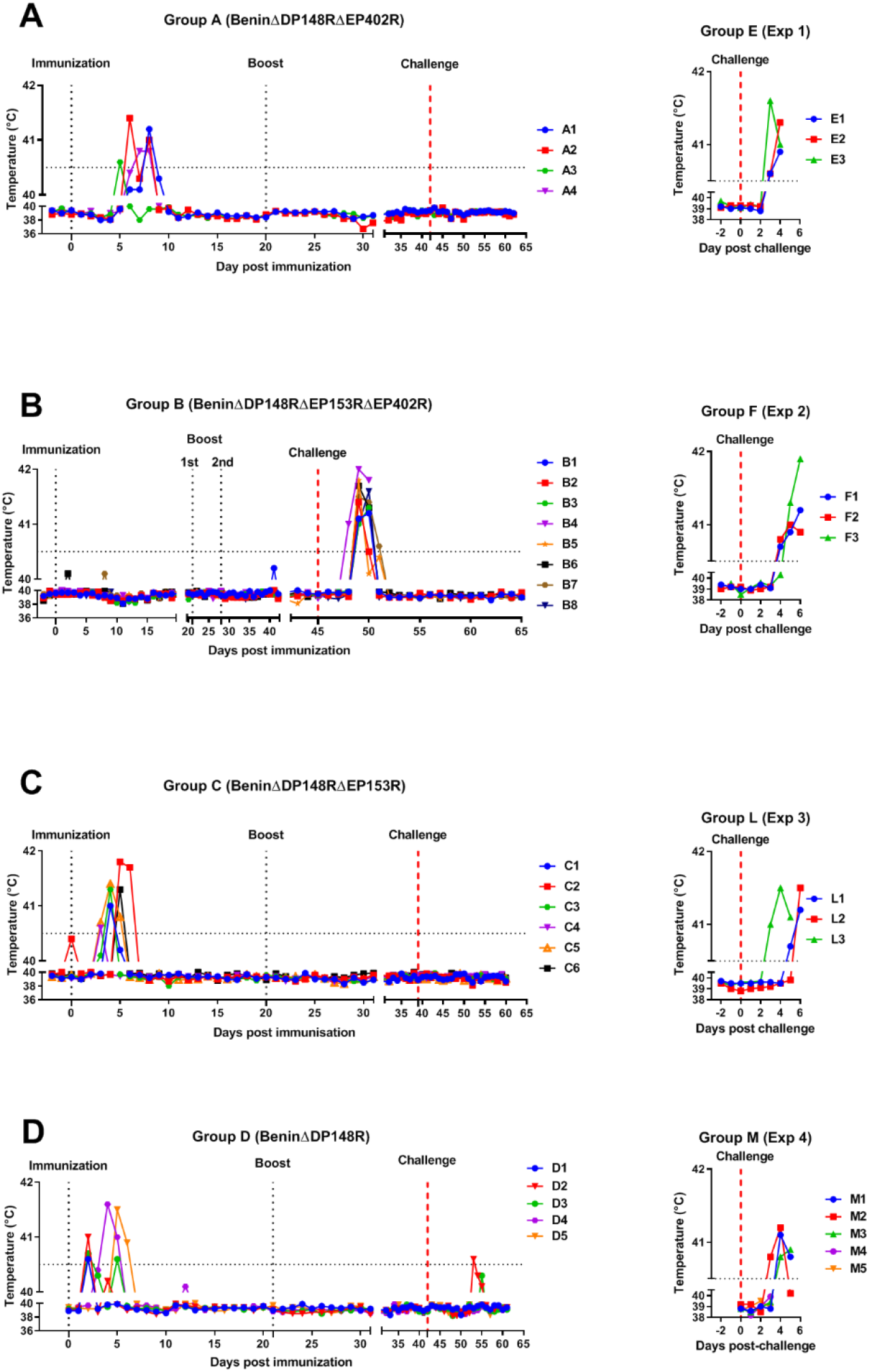
Temperatures following immunization and boost of pigs with BeninΔDP148RΔEP402R (Group A), BeninΔDP148RΔEP402RΔEP153R (Group B), BeninΔDP148RΔEP153R (Group C) and with BeninΔDP148R (Group D) and challenge with Benin 97/1. Temperatures for non-immune control pigs after challenge (Group E, Group F, Group L and Group M).

**Figure 5.**
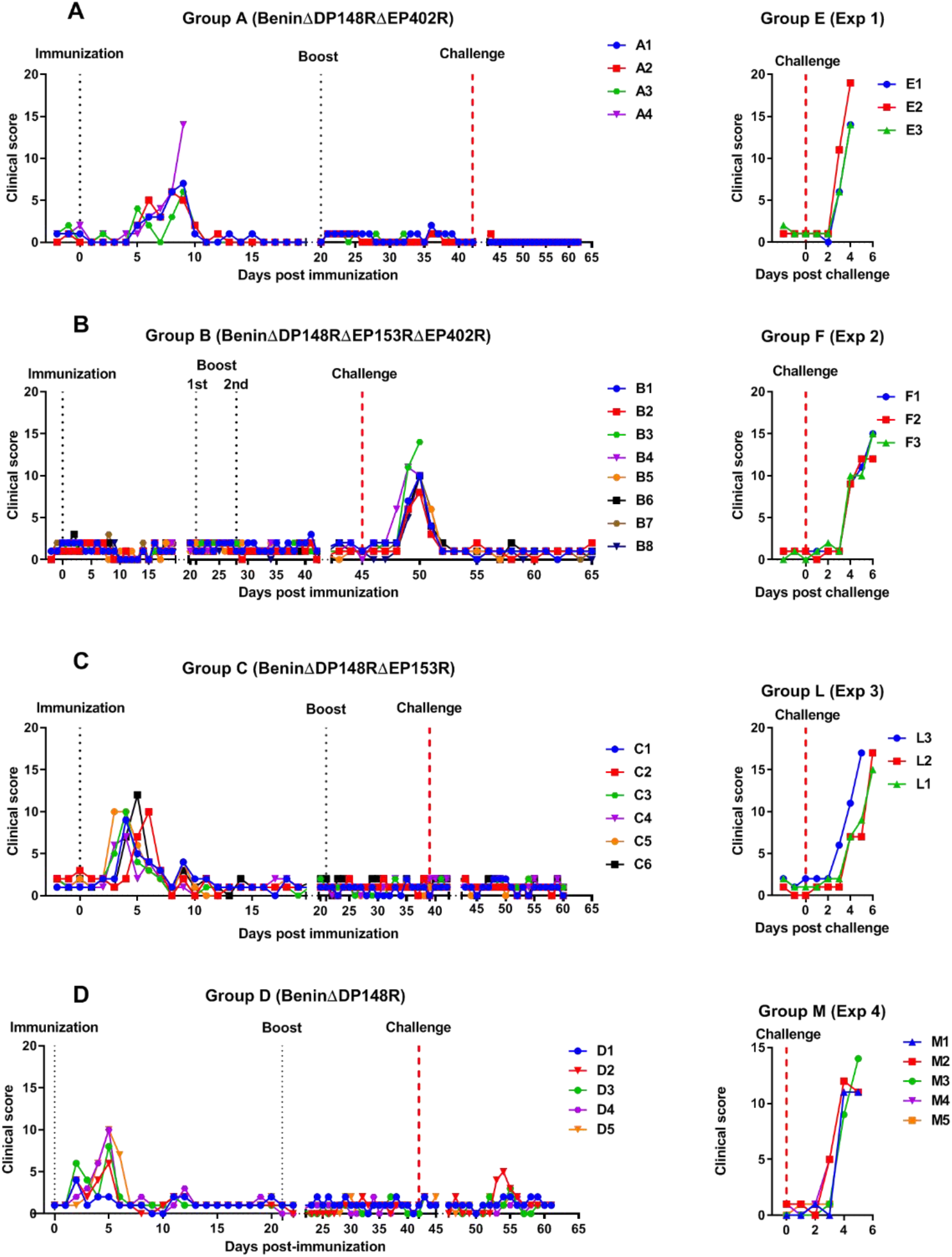
Clinical scores following immunization and boost of pigs with BeninΔDP148RΔEP402R (Group A), BeninΔDP148RΔEP402RΔEP153R (Group B), BeninΔDP148RΔEP153R (Group C) and with BeninΔDP148R (Group D) and challenge with Benin 97/1. Scores for non-immune control pigs after challenge (Group E, Group F, Group L and Group M).

In Group B (BeninΔDP148RΔEP153RΔEP402R) no increase in temperature or other clinical signs were observed in any of the pigs before challenge. After challenge an increase in temperature at or above 41°C was observed in one pig at 3 days post-challenge and in the remaining pigs at day 4 post-challenge (Figure 4B). Pigs B3 and B4 also had breathing difficulties on day 5 post-challenge, reaching the humane endpoint and were euthanized (Figure 5B). The remaining 4 pigs had an increased temperature above 40.5°C for two days in total, except for pig B7 which had a temperature of 40.6°C which persisted for 3 days (Figure 4B). Non-immune pigs in group F also developed clinical signs at between day 2- and 4 post-challenge, including an increase in temperatures (40.3 – 41.9°C), were not eating and were lethargic (Figures 4B and 5B). At day 6 post-challenge, Pig F2 had hemorrhagic lesions at the periphery of the ears while Pig F3 had traces of blood in its feces. On days 4 to 6 post-challenge all pigs were euthanized at the moderate severity humane endpoint.

In Group C (BeninΔDP148RΔEP153R), moderate clinical signs were observed after immunization, comparable with group D (BeninΔDP148R), with increased temperatures above 40.5°C for one or two days between days 3 to 6. However, after challenge none of the pigs from either of the groups showed any clinical signs or temperatures (Figures 4C, 4D, 5C and 5D). In clear contrast, control pig L3 health status deteriorated quickly starting from day 3 post challenge, with a raise in rectal temperature above 41.5°C at day 4 post challenge. Pig L3 was euthanized at day 5 post challenge and the rest of the group reached the humane end point at day 6 post challenge due to increased temperatures (41.2 – 41.5 °C) and rapid manifestation of clinical signs including lethargy and refusal to eat (Figures 4C and 5C).

In group D (BeninΔDP148R), pigs D1, D2 and D3 had increased temperatures at day 2 post-immunization, with temperatures ranging from 40.6 to 41°C. Pigs D4 and D5 showed increased temperatures on day 4, but these dropped on the following day (Figure 4D). No temperatures or clinical signs were present after boost or challenge in this group (Figures 4D and 5D). Control pigs from group M showed clinical signs including increased temperature lethargy and reduced appetite or refusal to eat from day 2 or 3 post challenge all were euthanized at day 4 or 5 post challenge (Figure 4D and 5D).

### Post mortem pathological observations

Pigs in Group A, immunized with BeninΔDP148RΔEP402R, showed few macroscopic lesions, with the exception of the slight enlargement of lymph nodes. Pig A4, was euthanized due to welfare reasons before challenge (Figure 6). In Group B, immunized with BeninΔDP148RΔEP153RΔEP402R, the pigs culled at humane end points (B3, B4) had higher macroscopic lesion scores than those which survived but lower than the control pigs. Pigs B3 and B4 had ascites, moderate/partial hyperemic splenomegaly, several enlarged lymph nodes and hemorrhagic renal lymph nodes. Three surviving pigs (B1, B7 and B8) had enlarged lymph nodes while the other two (B2, B5) were free of any ASFV typical lesions (Figure 6). In group C, immunized with BeninΔDP148RΔEP153R, pigs C1 and C2 showed enlarged spleens and minimal hemorrhages on the lymph nodes (renal and gastro hepatic) (Figure 6). Similar findings were reported in group D (BeninΔDP148R), where pigs D3, D4, D5 had enlarged lymph nodes, and pig D2 had an enlarged spleen and pericardial effusion (Figure 6). All control pigs belonging to groups E, F, L and M presented lesions consistent with acute ASF, represented by enlarged and hemorrhagic lymph nodes, erythematous tonsils, pericardial effusions, enlarged spleens, and ascites.

**Figure 6.**
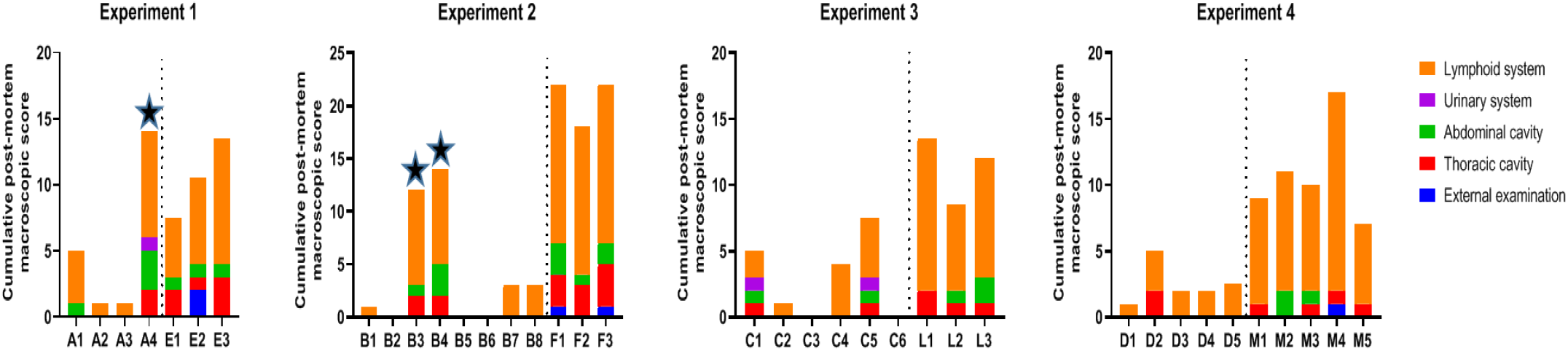
Scoring macroscopic lesions at the cull point for the vaccinated groups: Experiment 1, BeninΔDP148RΔEP402R (Group A), Experiment 2, BeninΔDP148RΔEP402RΔEP153R (Group B), Experiment 3, BeninΔDP148RΔEP153R (Group C) and Experiment 4, BeninΔDP148R (Group D) and the control groups (E, F, L and M). Lesions are presented on the graph by different colours. Stars represent the animals that reached the end-point before study termination.

### Genome copies in blood

Levels of virus genome in blood were measured by qPCR. At 5- or 6-days post-immunization 3 pigs in Group A, (BeninΔDP148RΔEP402R) had moderate levels of virus in blood (approximately 10^6^ genome copies/ml) coinciding with the onset of clinical signs, whereas pig A3 only had low levels of virus (approximatively 10^3^ genome copies/ml) (Figure 7A). Virus levels decreased after day 6 post-immunization and no pigs had detectable virus after boost and challenge.

**Figure 7.**
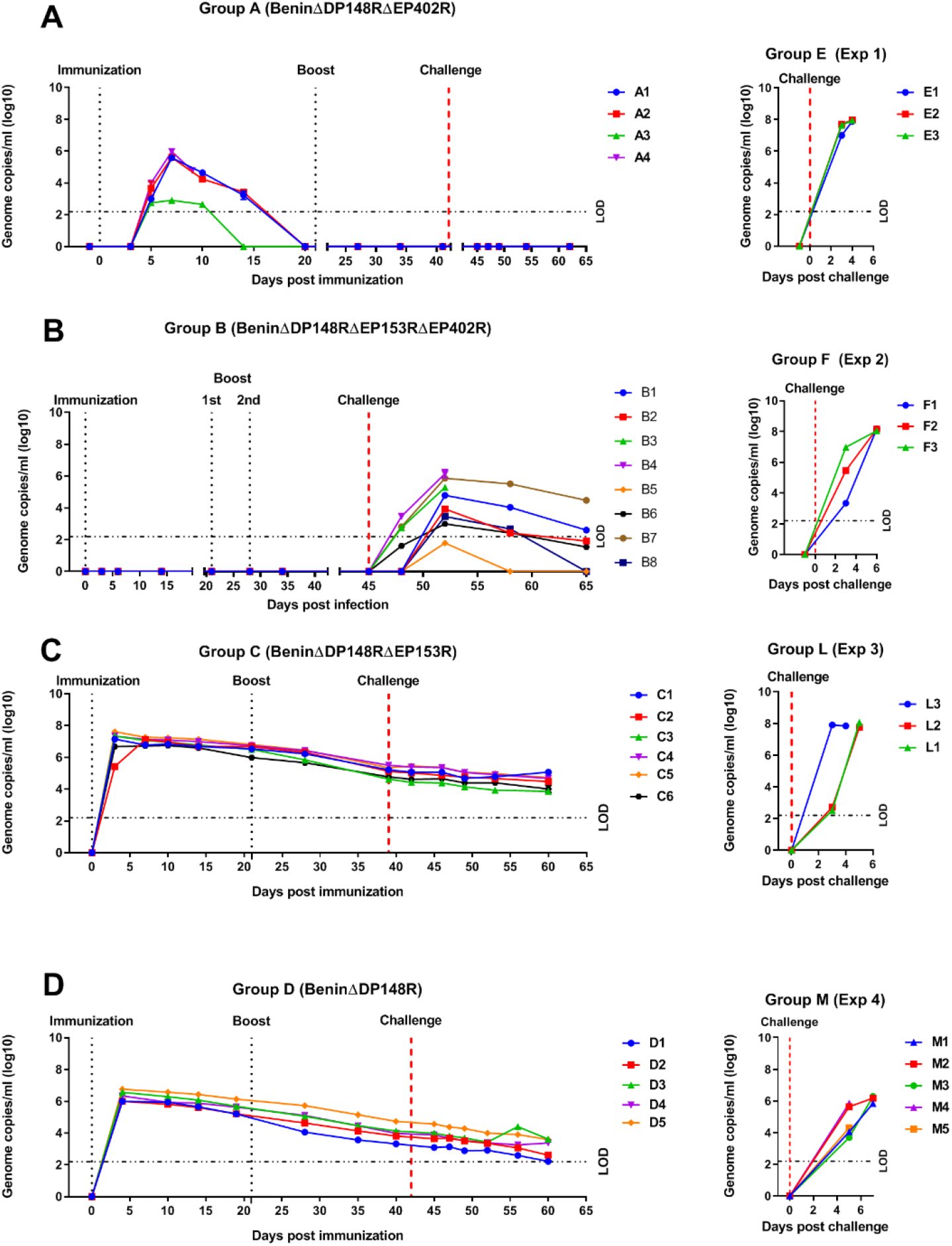
Levels of viral genome in blood of pigs immunized with BeninΔDP148RΔEP402R (Group A), BeninΔDP148RΔEP402RΔEP153R (Group B), BeninΔDP148RΔEP153R (Group C), BeninΔDP148R (Group D). Levels of control groups corresponding to each study are presented in the right panels (Groups E, F, L, M). Results are estimated by qPCR and reported as genomic copies/ml (log10) of blood. Dashed line represents the limit of detection (LOD) of the assay.

No virus genome was detected in blood from pigs in Group B (BeninΔDP148RΔEP153RΔEP402R) before challenge (Figure 7B). After challenge variable levels of genome were detected from day 3 or 4 post-challenge. Maximum levels were detected in pigs B3 and B4, which reached the humane endpoint (approx. 10^6^ genome copies per ml). In the surviving 6 pigs the maximum levels detected varied between ∼10^2^ and 10^5.5^ genome copies/ml. Levels of virus genome decreased until termination at day 20 post-challenge becoming undetectable in 2 pigs and reduced to 10^2.6^ in others except 1 pig were levels remained at 10^4.5^ at termination.

Pigs belonging to groups C (BeninΔDP148RΔEP153R) and D (BeninΔDP148R) had higher levels of genome copies in blood (approx. 10^6^-10^7^ genome copies/ml) following immunization, these levels decreased to approx. 10^4^ genome copies/ml by the end of the study (day 60) (Figures 7C and 7D). Interestingly levels of copies in blood were significantly higher at day 10 post immunization (p<0.01) for group C compared with group D and this difference persisted throughout the study (p<0.1 at days 14, 28, 39/40 and 49 post immunization) (Figure 8).

**Figure 8.**
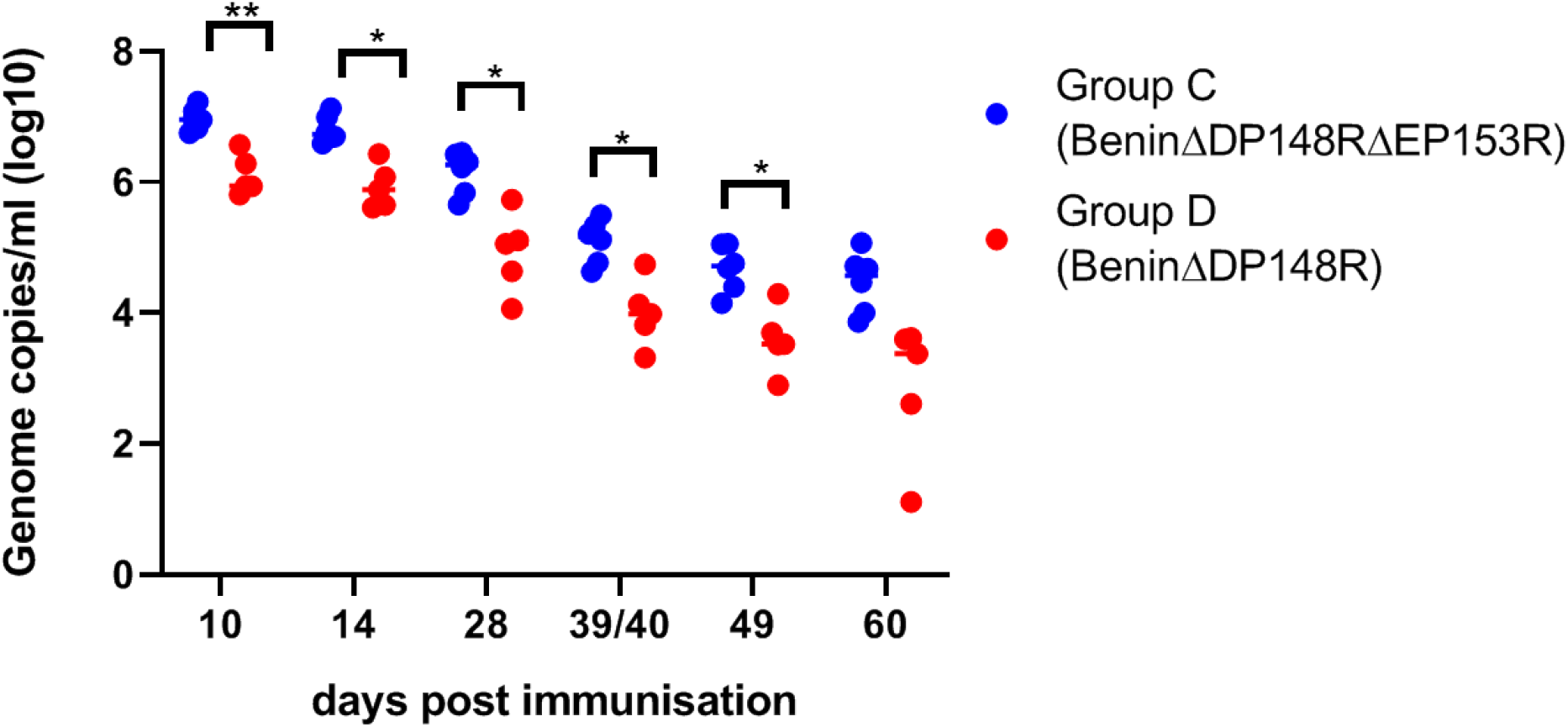
Levels of viral genome in blood of pigs immunized with BeninΔDP148RΔEP153R (Group C) and BeninΔDP148R (Group D). Statistically significant responses between groups are presented (** is p<0.01; * is p<0.1)

### IFN-gamma ELISpot assay

The responses of PBMCs from immunized pigs to ASFV were measured at different times post-immunization by IFN-γ ELISpot. An ASFV specific response was not detected before immunization in any group of pigs. Very high numbers of IFN-γ producing cells (∼725 – 1225 spots per million cells) were induced in all pigs in Group A before boost, which then decreased before challenge to levels ranging from 178 - 463 spots/million cells (Figure 9A). In contrast, for Group B, low levels of IFN-γ producing cells were induced after immunization and before the first boost. Hence, a second boost with higher dose of the same virus was applied 4 weeks post immunization and the number of IFN-γ producing cells increased uniformly in all pigs before challenge (326 – 540 spots/million PBMCs) (Figure 9B). Of note, pigs B3 and B4 that were euthanized at 5 dpc had 540 and 458 spots/million PBMC. Pigs in group C showed a relatively lower number of IFN γ producing cells before boost, with pig C2 showing a higher response of ∼305 spots/million PBMCs, compared with other pigs of this group. Surprisingly, levels decreased after the boost and only one pig (C5) had a higher number of IFN-γ secreting cells (135 – 126 spots/million PBMCs) at challenge compared with before boost (Figure 9C). The number of IFN-γ producing cells were significantly higher in group A compared with groups B and C before boost. At challenge, both groups A and B showed significantly higher number of IFN-γ producing cells than Group C (Figure 9D).

**Figure 9.**
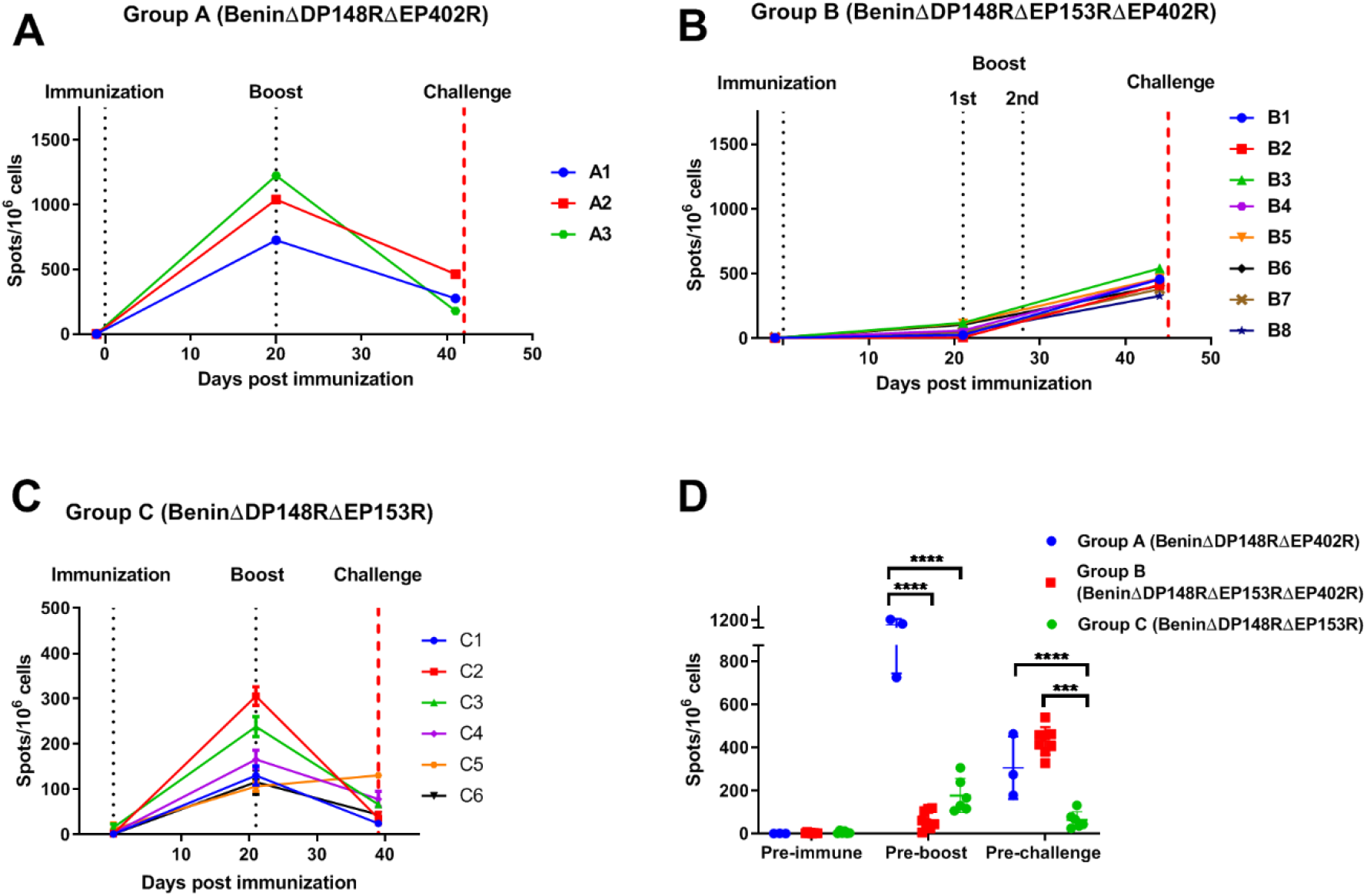
IFNγ responses from PBMCs collected post immunization, post boost and before challenge. Panels A and B show the number of IFNγ secreting cells stimulated with Benin 97/1 by ELISPOT assay. PBMCs were collected from pigs immunized with BeninΔDP148RΔEP402R (Group A), BeninΔDP148RΔEP402RΔEP153R (Group B), and BeninΔDP148RΔEP153R (Group C). Results are presented as mean number of IFNγ producing cells/10 6 cells. Statistically significant responses between groups pre-boost and pre-challenge are presented in Panel D (**** is p<0.0001; *** is p<0.001).

### Antibody responses

Antibody responses to ASFV p72 capsid protein were measured using a commercially available blocking ELISA. As expected, an antibody response was detected in all pigs in Group A by day 14 and increased after boost (Figure 10A). In contrast, pigs in Group B mounted a slower antibody response, which was first detected at day 27 post-immunization after the first boost and just before the second boost. Levels were maintained for the rest of the experiment (Figure 10B). A faster antibody response was seen in group C, where most of the pigs had detectable antibody by day 7, and on day 14 all were above the cut-off (Figure 10C). For group D, a general trend was observed in generating an antibody response at day 10 (Figure 10D).

**Figure 10.**
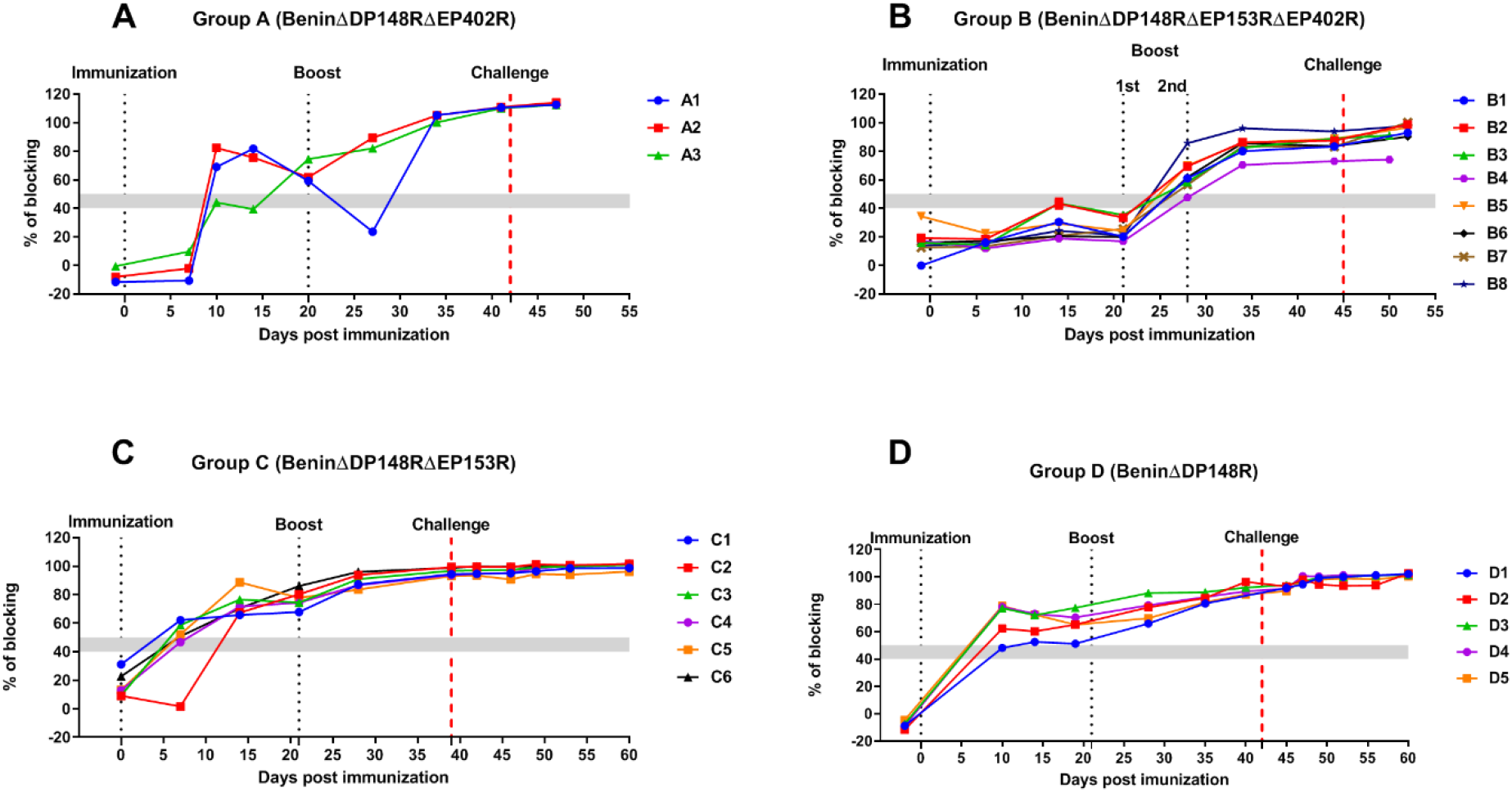
Antibody responses post immunization. Serum samples were collected from pigs immunized with BeninΔDP148RΔEP402R (Group A), BeninΔDP148RΔEP402RΔEP153R (Group B), BeninΔDP148RΔEP153R (Group C) and BeninΔDP148R (Group D) and assayed using a commercial blocking ELISA against p72 protein. Results are presented as percentage of blocking and cut-off value is represented by 50%.

## Discussion

Targeted deletions of non-essential genes from the ASFV genome have been used successfully to construct candidate live attenuated vaccine strains and to understand the role of the genes during infection of cells and pigs. A vaccine candidate with an acceptable safety profile should have reduced clinical signs and vaccine virus persistence post-immunization but retain high levels of protection.

One of the potential candidates for targeted deletions represents the EP402R gene which codes for the CD2v protein, since it is required for the binding of infected cells or viral particles to red blood cells, thus playing an important role in viral dissemination in different tissues in pigs. During the sylvatic cycle, in ticks, the binding of the virus to red blood cells probably helps retention of the virus with the blood meal and also crossing the midgut barrier (35). Deletion of the EP402R gene from the genome of virulent viruses Malawi Lil20/1 (genotype VIII) (16) and Georgia/07 (genotype II) (12) did not reduce the virulence of these viruses in pigs. Surprisingly, deletion of this gene from a genotype I virulent isolate, BA71, reduced the virus virulence and induced protection against lethal challenge with the parental virus and also against the genotype II Georgia/07 isolate (36). Recent studies have shown that deletion of an additional virulence marker such as UK (DP96R) gene resulted attenuation in pigs, conferring 100% protection after challenge with an Asian strain belonging to genotype II (ASFV-SY18), however viral DNA was still present in different lymphoid organs (37). Deletion of the EP153R gene from the genome of a virulent isolate, Malawi Lil20/1, failed to reduce virulence of the virus for pigs (38). Both the EP402R and EP153R genes are interrupted in the genomes of the naturally attenuated isolates OURT88/3 and NH/P68 (26, 39). However, the role of these gene interruptions in virus virulence is unclear since OURT88/3 and NH/P68 viruses also have large deletion of multiple genes belonging to MGF 360 and MGF 505 from close to the left genome end (26, 40). Both the single deletion of EP402R and the simultaneous deletion of EP402R and EP153R from an attenuated Georgia/07 strain (ASFV-G-Δ9GL) decreased the ability to protect against challenge (41). Serum antibodies against both CD2v and EP153R were shown to mediate haemadsorption inhibition and to be involved in serotype-specific protective immunity, representing good targets for ASFV serotype classification and evolution (42, 43).

Our approach to reduce the virus persistence and clinical signs post-immunization of pigs with BeninΔDP148R (7), was to remove either EP402R or EP153R genes singly, or in combination from the BeninΔDP148R attenuated virus. Our results showed that immunization of pigs with the virus BeninΔDP148RΔEP402R resulted in a dramatically shorter virus persistence compared to BeninΔDP148R, with virus genome in blood detected only until day 14 post-immunization compared with more than 60 days in pigs immunized with BeninΔDP148R (Figure 7A and 7D). Moreover, no viremia was detected after the second boost or after challenge, confirming our hypothesis that CD2v can play a role in viral persistence. However, since the immunized pigs still displayed significant clinical signs post immunization, it is also an indicator that the virus might be disseminated by another route.

We were surprised that deleting the EP402R gene did not have a greater effect on reducing clinical signs post-immunization since attenuation was observed when this gene was singly deleted from the virulent BA71 virus, another Genotype I isolate (36). In order to try to further increase the safety profile of BeninΔDP148RΔEP402R, we decided to also remove the EP153R gene, which is adjacent to EP402R (26). This gene codes for a type II membrane protein containing a C-type lectin domain similar to those in host proteins. The EP153R protein has been shown to have diverse roles, including increasing the binding of red blood cells to ASFV infected cells, inhibition of cell surface expression of SLA-I (porcine MHC class I) and inhibition of apoptosis (23-25). It was also shown that together with CD2v, EP153R can contribute to mediating the cross-protective serotype specific immunity (43).

After immunization with BeninΔDP148RΔEP153RΔEP402R (Group B), pigs did not show any clinical signs or viremia before challenge (Figures 4B, 5B, 7B). However, when we deleted EP402R or EP153R singly, elevated clinical signs were observed after immunization. This could be explained if the proteins act synergistically, thus the deletion of both genes reduces the viral load to an extent that the innate immune response can suppress the initial steps of the viral replication. A faster antibody response was mounted in groups A (BeninΔDP148RΔEP402R), C (BeninΔDP148RΔEP153R) and D (BeninΔDP148RΔEP153R), whereas group B (BeninΔDP148RΔEP153RΔEP402R) showed a slower antibody response, since the animals seroconverted only after the second boost (Figure 10). When looking at the cell-mediated immune responses, PBMCs collected from Group A (BeninΔDP148RΔEP402R) had higher responses than Group B (BeninΔDP148RΔEP402RΔEP153R) and Group C (BeninΔDP148RΔEP153R) before the boost (Figure 9D). Interestingly, PBMCs from pigs immunized with BeninΔDP148RΔEP153RΔEP402R only had a considerable response after boost with numbers increasing before challenge (Figure 9B) and no statistical significance was observed between Groups A and B at that time (Figure 9D). These results strengthen our hypothesis of a synergistic effect of deleting EP153R and EP402R genes in reducing viral replication and persistence. This is supported by the fact that the C-type lectins are reported to be involved in the interaction with glycans on the cells surface and may facilitate the attachment of the viruses to different cells.

After challenge, Group B (BeninΔDP148RΔEP153RΔEP402R) presented moderate viremia (up to 10^6^ genome copies/ml) (Figure 7B), with 2 pigs reaching the endpoint, indicating that the immune response induced was not sufficient to suppress replication of the challenge virus. As discussed above, possibly BeninΔDP148RΔEP402RΔEP153R did not replicate to the same extent, as no viremia was detected, and consequently a reduced immune response was induced. Since all viruses replicated to a similar level as parental virus in macrophages *in vitro*, the reduced replication *in vivo* probably resulted from interactions with other host cells. Group C (BeninΔDP148RΔEP153R) showed a slightly higher viremia level when compared with Group D (BeninΔDP148R) (Figure 8). These differences were observed as early as day 10 post immunization, suggesting that they were not mediated by specific antibody or T cell responses against EP153R in animals infected with BeninΔDP148R.

In summary, we showed that by removing EP402R from the BeninΔDP148R backbone we reduced viral persistence in blood after immunization, reducing clinical signs but maintaining a survival rate of 100% after challenge and no replication of challenge virus. More importantly, the additional deletion of EP153R increased attenuation, since no clinical signs or viremia were observed after immunization, however elevated clinical signs and viremia were observed after challenge with 75% survival rate. Deletion of EP153R alone did not reduce virus persistence or clinical signs after immunization compared to the single deletion of DP148R gene. These results highlight an important role for both EP153R and EP402R proteins in promoting virus replication *in vivo*. This may be mediated by the cooperation of both proteins in binding of virus particles and infected cells to red blood cells. In addition, the two proteins may also act singly or synergistically to evade initial steps of the immune response and a role in immune modulation has already been shown for the EP402R protein. C-type lectin proteins have diverse roles in mediating cell to cell adhesion. For example, C-type lectin receptors play a crucial role in Natural Killer (NK) cell activity. It is tempting to speculate that EP153R may have evolved to evade NK host immune responses since NK activity was shown to correlate with protection following immunization with the attenuated NH/P68 isolate (40).

Overall, the viruses we have constructed from already attenuated virus provide the means to dissect the role of these and other viral proteins in pathogenesis and evasion of immune responses.

## Acknowledgements

We are grateful to Animal Services at Pirbright for help with animal experiments and members of the ASFV team for helpful discussions.

